# Engineering a membrane-independent human prothrombinase through parsimonious mutation of factor Xa

**DOI:** 10.1101/2025.04.16.649061

**Authors:** Fatma Işık Üstok, James A. Huntington

## Abstract

**Background:** Thrombin is generated from its precursor prothrombin by sequential cleavage at Arg320 and Arg271 by the prothrombinase complex, composed of factor (f) Xa and fVa on phospholipid (PL) membrane surfaces. The affinity of human fXa for fVa is low in the absence of PL. However, fXa orthologues from the venom of group D snakes bind to fVa with high affinity, forming an active complex without PL. We previously characterized the properties of the fXa orthologue Hopsarin D (HopD) from *Hoplocephalus stephensii*.

**Objectives:** Here we set out to create a PL-independent human prothrombinase by making mutations to fXa, guided by a model of the prothrombinase complex and the sequence differences between human fXa and HopD.

**Methods:** We assessed the contribution of individual domains of fXa to its binding to fVa by swapping each with the corresponding domain of HopD. We then chose three loops in the serine protease (SP) domain predicted to be in contact with fVa to swap to those of HopD. Eventually, 10 residues from the three loops in the SP domain and 7 from the EGF2 domain were selected for mutation.

**Results:** The resulting ‘M17’ fXa variant bound to fVa with a Kd of ~20 nM, similar to HopD, and together efficiently processed prothrombin through the meizothrombin intermediate in the absence of PL.

**Conclusions:** We conclude that the role of PL membranes in prothrombinase assembly and function is limited to improving the affinity of fXa for fVa. The M17-fVa complex is likely to be structurally equivalent to the human prothrombinase complex.

## INTRODUCTION

Thrombin is generated from its circulating precursor prothrombin through cleavage of two bonds by the enzyme complex known as prothrombinase [1]. Assembly of the protein components of prothrombinase, the cofactor factor (f) Va and the serine protease fXa, is critically dependent on the presence of Ca^2+^ and a negatively-charged phospholipid (PL) membrane surface [2–4]. Factor Va is a two-chain molecule after activation by fXa or thrombin, and is comprised of three A domains and two membrane-anchoring C domains; the heavy chain contains the A1 and A2 domains, and the light chain the A3, C1 and C2 domains (Figure 1A). Factor Xa is also a two chain molecule comprised of a membrane-binding gamma carboxyglutamic acid (Gla) domain and two epidermal growth factor-like (EGF) domains (light chain), linked via a disulfide bond to the serine protease (SP) domain (heavy chain; Figure 1B). The PL-surface improves the affinity of the components by about 3-orders of magnitude, primarily through co-localization and reduction in diffusional dimensionality [5]. Some studies have suggested that conformational change in fXa upon PL binding also plays a role in enhancing binding to fVa, presumably by altering the interactions between the N-terminal Gla and EGF1 domains with the C-terminal EGF2 and SP domains [6–8]. The substrate prothrombin also interacts with the PL surface, resulting in a modest increase in rate of thrombin generation [9]. It has also been reported that the interaction of prothrombin with the PL surface is critical for presenting Arg320 for initial cleavage by prothrombinase [9,10]. The PL membrane surface therefore plays a critical role in prothrombinase assembly and function, with PL-dependence limiting thrombin generation to sites of tissue damage. Snakes of the Elapidae family have evolved venom orthologues of the components of prothrombinase that operate in the absence of PL membranes to kill their prey by triggering thrombin generation throughout the vasculature, leading to micro-thrombosis in organs and hemorrhage through consumption of coagulation proteins and platelets [11]. We previously characterized a fXa venom orthologue Hopsarin D (HopD) that binds to human fVa with high affinity (Kd of 10 nM) and activates prothrombin efficiently in the absence of PL membranes [12]. HopD with fVa processes prothrombin in a manner similar to fully assembled human prothrombinase, with initial cleavage at Arg320 to form the active intermediate meizothrombin, followed by cleavage at Arg271 to release thrombin. This work supports the contention that the role of the PL membrane is primarily to help assemble prothrombinase, and is not required to enforce the order of prothrombin cleavage. However, HopD is not human fXa and contains insertion loops that might influence the binding to and processing of prothrombin.

**Figure 1.**
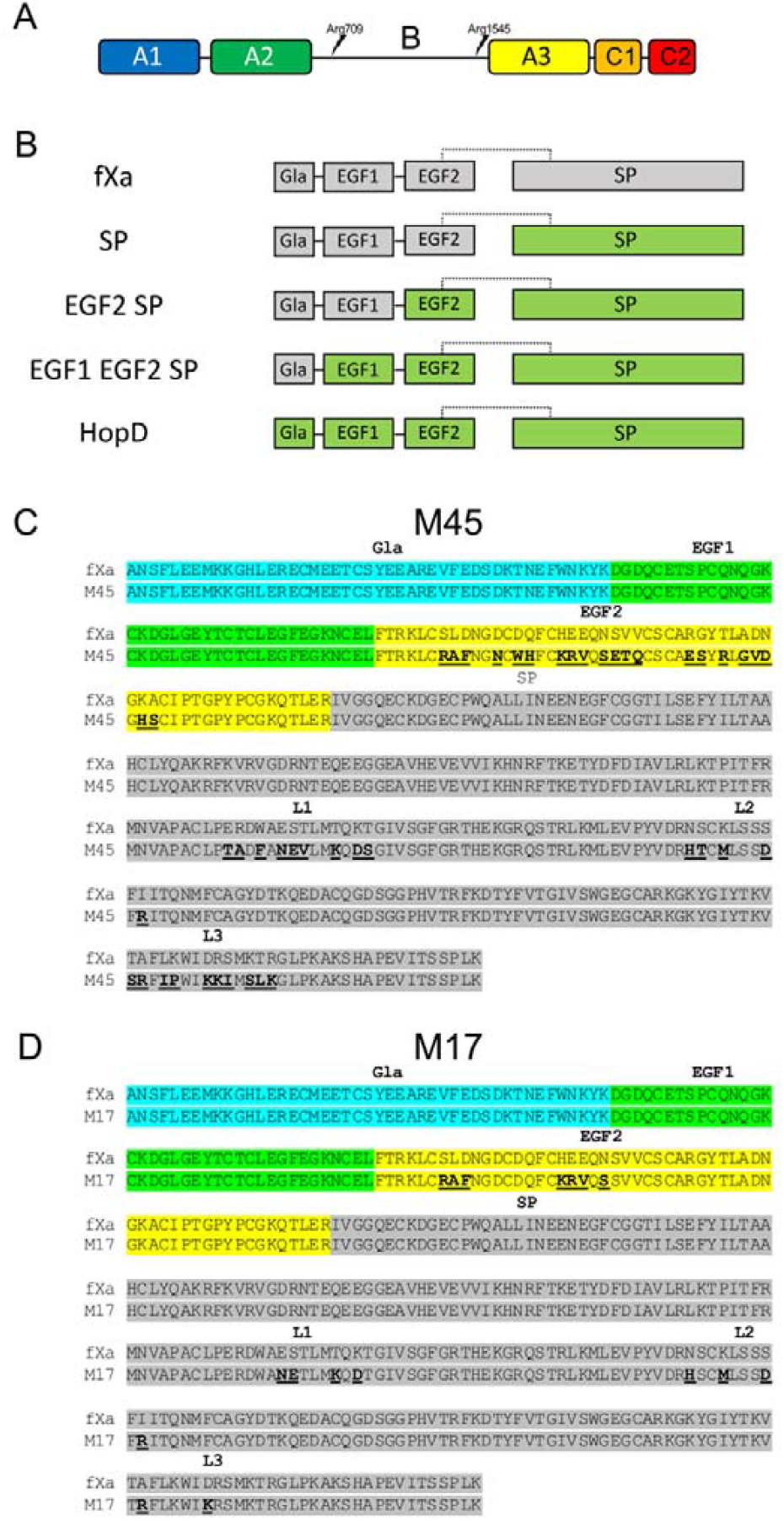
Factor V, fXa and HopD domain organization, chimeric constructs and sequences alignments. (A) Schematic of domain organization of fV, colored from N-terminus (blue) to C-terminus (red). Idividual domains are denoted, with the A1 and A2 domain constituting the heavy chain and the A3 and C domains the light chain, after activation by excision of the B domain by proteolysis at sites indicated. (B) Factor Xa (fXa; gray) and Hopsarin D (HopD; green) share the domain organization of N-terminal gamma carboxyglutamic acid (Gla) domain and two epidermal growth factor-like (EGF) domains, linked via disulfide bond (dashed line) to a serine protease (SP) domain. These domains were swapped sequentially from the C-terminal SP domain to assess the contributions towards binding affinity to fVa. (C) Sequence alignment of wild-type fXa and the EGF2/L123 chimera (M45), with the Gla, EGF1, EGF2 and SP domains highighted in cyan, green, yellow and grey, respectively, with the L1, L2 and L3 regions indicated. The residues with sequence differences relative to human fXa are in bold and are underlined. (D) Sequence alignment of fXa and the M17 variant, as in C.

In order to investigate the PL-dependence of prothrombinase assembly and function, we set out to create a membrane-independent human fXa with mutations guided by sequence differences with HopD and a molecular model of human prothrombinase [13] based on the crystal structure of the venom prothrombinase pseutarin C [14]. We succeeded in producing a variant with only 17 mutations (M17) that bound to fVa with high affinity and processed prothrombin down the meizothrombin pathway in the absence of PL membranes. The properties of M17 have implications for the assembly and function of human prothrombinase, and M17 may also be a useful tool for use in structural studies, eliminating the requirement for added PL.

## METHODS

### Chimera construct design

Primers used to introduce domain swaps and point mutations had sequence-specific overlapping regions to enable correct assembly using the Klenow-based ligation-independent assembly method [15]. Fragments were amplified by PCR from a previously generated pCEP4-human fX plasmid using appropriate primers. All PCR procedures were conducted using Q5 Hot Start High-Fidelity 2X Master Mix (New England Biolabs) according the manufacturer’s instructions. After the PCR reaction, each fragment was purified, Dpn1 digested, and gel extracted. The plasmid backbone was isolated by restriction digestion of a previously constructed pCEP4-fX vector with Kpn1/EcoRV restriction enzymes, with a band (9149bp) containing the plasmid backbone excised from an agarose gel, purified and used in the assembly procedure. The purified fragments and plasmid backbones were added to the Klenow assembly mixture in equimolar amounts, incubated at 37°C for 20 min, followed by cooling on ice and transformation into competent *E.coli* cells (Max Efficiency DH5α competent cells, Invitrogen). The same method was used to generate individual and combined chimeric variants as well as point mutations. All of these constructs contained the prothrombin pro- and signal regions to enable gamma-carboxylation [16]. The chimeric fX constructs containing entire HopD domains were generated using the same assembly method as described above. The same method was used to generate truncated M17, where the pCEP4-M17 FL plasmid was used to amplify EGF2 and SP domains (regions 85 to 425, mature numbering) and cloned into pET23 vector. All plasmid sequences were confirmed by DNA sequencing prior to use in maxi-prep preparations for HEK-EBNA cell transfections to generate stable cell lines.

### Expression and purification of recombinant proteins

HEK-EBNA cells were transfected with the fX constructs and selection, purification and activation were conducted essentially as previously described [12,17]. Briefly, cells expressing fX were grown in CD-CHO media supplemented with 4 mM L-glutamine, geneticin (G418, 0.025 mg/ml), hygromycin B (200 μg/ml) and vitamin K1 (10 μg/ml). Media was harvested every 2 to 3 days, pooled and concentrated 10-fold using a Vivaflow 200 (Sartorius) concentrator. Trisodium citrate was added to the concentrated media to a final concentration of 15 mM, followed by stirring for 30 minutes at 4°C. Iced barium chloride (1M stock; 80 ml/L media) was added slowly with stirring to precipitate proteins containing gamma-carboxylated Gla domains, followed by 30 minutes with stirring and 1h stationary incubations at 4°C. Precipitate was pelleted by centrifugation at 6000 x g for 15 minutes at 4°C. The recovered pellets were dissolved in 0.2 M EDTA (150 ml/L media), followed by dialysis against 5 L 20 mM Tris pH 7.4, 50 mM NaCl, 1 mM EDTA and 1 mM benzamidine-HCl overnight at 4°C. Dialysates were filtered before loading onto a HiTrap Q Sepharose column (5ml, Cytiva) equilibrated with 20 mM Tris pH 7.4, 50 mM NaCl, and protein was eluted with a gradient from 0.05-0.7 M NaCl over 7 column volumes (CVs). The recovered protein was then diluted 10-fold in 20 mM imidazole pH 6.5, 5 mM CaCl_2_ and loaded onto a heparin Sepharose column (5 ml, Cytiva) equilibrated with the same buffer. The protein was eluted with a gradient of 0-0.75 M NaCl over 10 CVs. A sample (5 μl) of each fraction was pipetted into a well of a 96-well plate containing 45 μl Tris-Buffered Saline (TBS) with 5 mM CaCl_2_ containing Russell’s viper venom factor X activator (RVV-X, 1:100 w/w) and chromogenic substrate S-2222 to determine the fractions containing factor Xa activity. Fractions with activity were pooled and concentrated for further purification on a Superdex 200 (PG 16/600, Cytiva) column equilibrated with 20 mM Tris pH 7.4, 150 mM NaCl and 5 mM CaCl_2_. The purified fX was activated with RVV-X (1:100 w/w) by incubation at 37°C for 3 hours., followed by an additional purification step on heparin Sepharose column (5ml, Cytiva), as described above. Protein was concentrated to 2 mg/ml) and dialyzed into 20 mM Tris pH 7.4, 150 mM NaCl and 5 mM CaCl_2_ buffer. The activity of all fXa variants against S-2222 was assessed to ensure that proteolytic activity was not affected by the domain swaps and mutations (Supplemental Figure 1).

Stably transfected BHK-M cells were used to express fV_BD_ (B-domainless human fV), and purified protein was activated to fVa when required as described previously [18]. Recombinant human prothrombin (S195A) was prepared as described previously [17]. Truncated M17 (containing the EGF2 and SP domains) was expressed in *E.coli* (BL21 STAR DE3 pLysS) as inclusion bodies, refolded, purified and activated essentially as previously described [12,19].

Chymotrypsin template numbering is used throughout when referring to residues in the SP domain of fXa or its variants.

### Prothrombin processing monitored by SDS-PAGE

Prothrombin processing was monitored and evaluated as described previously [12]. Briefly, reactions were initiated by adding fXa (to 5 nM) to a solution containing 5 μM S195A prothrombin and 50 nM fV_BD_ (final concentrations) in assay buffer (20 mM Tris pH 7.4, 150 mM NaCl, 2 mM CaCl_2_, 0.1% PEG8000). Samples (10 μl) were withdrawn at indicated time points and quenched by mixing with 10 μl SDS sample buffer containing 100 mM DTT and 100 μM AEBSF, followed by incubation at 95°C for 5 minutes. Samples were run on 4-12% BisTris gradient gels (Invitrogen, Thermo Fisher Scientific) with MOPS buffer (Formedium). Bands were visualized by staining with Quick Coomassie Stain (Generon) at room temperature for 1 hour. Gels were destained and then imaged on a ChemiDoc MP Imaging System (Bio-Rad). Densitometry analysis was performed using Image Lab software (Bio-Rad) as described previously [12]. Two independent experiments were analysed separately for kinetics of appearance or disappearance of the reaction products using Prism (GraphPad, version 10.4.0), with one-phase decay for II consumption, one-phase association for H formation, and accumulation-decay for intermediates (F1.2-L and Pre-2). Rates given in the text are an average of two values with their standard error.

### Assessment of binding affinities of fXa variants to fV_BD_

Binding affinity was assessed using an Octet Red bio-layer interferometry (BLI) instrument, as described previously [12]. All experiments were performed in binding buffer (20 mM HEPES pH 7.4, 150 mM NaCl, 2 mM CaCl_2_, 0.1% BSA, 0.1% PEG8000) using Streptavidin-coated biosensors. fXa variants were treated with biotinylated EGRCK (EGR-chloromethyl ketone), desalted and loaded onto the biosensors at 1.0 μg/ml or 2.0 μg/ml. Remaining biotin binding sites on the biosensors were then quenched with biocytin (50 μg/ml). Recombinant fV_BD_ was diluted in binding buffer to concentrations between 4 nM and 5000 nM prior to the experiment. A no-analyte reference biosensor was used as a control for baseline drift. Data traces were analyzed using fortéBIO analysis software and Prism (GraphPad).

## RESULTS

### The contribution of individual domains to the affinity of fXa for fV_BD_

HopD and fXa share domain structure and considerable sequence identity. Removing the sequences of the activation peptide and the C-terminus (fXa has a long, unstructured C-terminus) allows for a more accurate assessment of the core sequence identity and similarity. 209 of the core 375 residues are identical between fXa and HopD (56%) and 52 additional non-identical residues have similar properties, resulting in a homology of 70%. HopD has two insertion loops relative to fXa in the SP domain, 2 residues after position 81, and 9 residues after position 90 (chymotrypsin numbering), that when removed from consideration improves identity and homology to 57% and 71%, respectively. The percent identity for the individual domains are: Gla 64% (28 of 44); EGF1 68% (25 of 37); EGF2 39% (18 of 46); and SP 57% (138 of 243). The EGF2 domain stands out with its relatively low sequence identity. To assess the importance of individual domains in contributing to the affinity of fXa for fVa, we swapped each domain from HopD into fXa sequentially.

Wild-type fXa and HopD were compared to chimeras where domains were swapped from HopD into fXa, starting with the C-terminal SP domain. The constructs are depicted in Figure 1B. Affinity was assessed by biolayer interferometry (BLI) with biotinylated fXa fixed to the tip of the probe and B-domainless fV (fV_BD_) used as the analyte. B-domainless fV is functionally similar to fVa [18] but less sticky, so the response vs. concentration curves fit nicely to a one-site specific binding model for assessment of binding affinity. Since determining relative binding affinities was our goal at this stage, binding experiments were conducted only once and on the same day. The Kds reported for the chimeras should therefore be considered as estimates, and were only used to guide the mutagenesis strategy. Wild-type fXa bound to fV_BD_ with a Kd of 1.4 μM and to HopD with a Kd of 18 nM, similar to the values obtained previously [12] (Figure 2A and 2E). Swapping of the entire HopD SP domain into fXa improved the binding affinity by 3.2-fold to a Kd of 400 nM (Figure 2B). Surprisingly, the addition of EGF2 to the SP chimera improved affinity an additional 15-fold to a Kd of 28 nM (Figure 2C). Adding EGF1 from HopD to the EGF2-SP chimera had little effect (1.3-fold; Kd of 21 nM; Fig. 2D), as did the addition of the Gla domain (1.2-fold; Kd of 18 nM). The difference in Gibbs free energy of binding between fXa and HopD is −2.58 kcal/mol, 90% of which is provided by the EGF2 and SP domains. Efforts to improve the binding of fXa for fVa would therefore focus on these two domains in which 133 residues are different between fXa and HopD.

**Figure 2.**
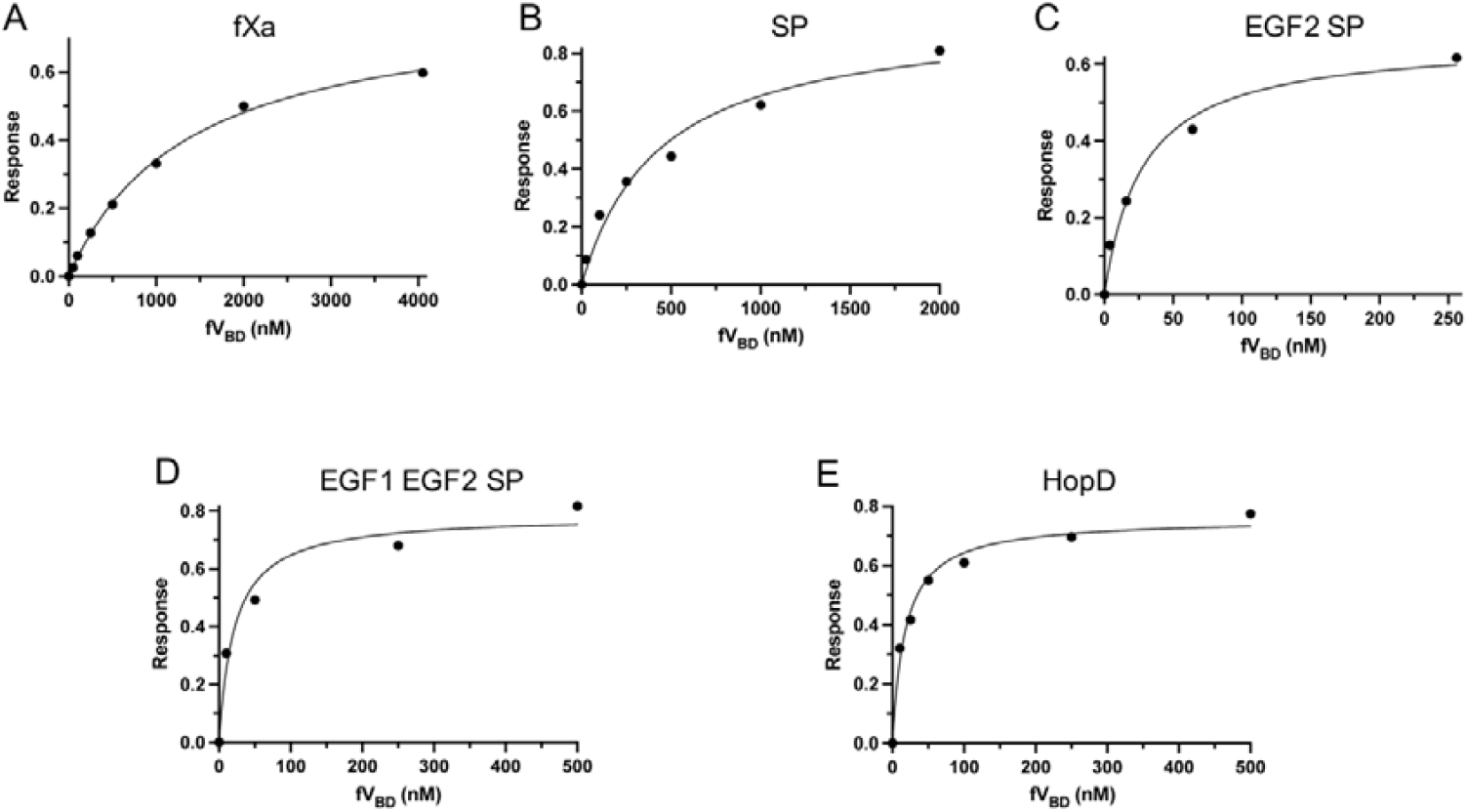
Binding of fXa domain-swap chimeras to fV_BD_ monitored by bio-layer interferometry (BLI). BLI response plotted against concentration of fV_BD_ is fit to a single site binding model to provide an estimate of the dissociation constant (Kd) for fXa (A), SP chimera (B), EGF2 and SP chimera (C), EGF1, EGF2 and SP chimera (D) and HopD (E).

### The contribution of individual SP loops to the affinity of fXa for fV_BD_

The model of human prothrombinase [13] was examined to identify candidate regions in the SP domain predicted to be in contact with fVa to swap out for the sequence of HopD. We chose three stretches roughly centered on the 130-helix (L1), the 170-helix (L2), and the C-terminal helix (L3). L1 is predicted to bind at the gap between the A2 and A3 domains of fVa, and to interact with the beginning of the hirudin-like a2-loop [20] at the C-terminus of the A2 domain. L1 is also at the interface with L3 and the EGF2 domain of fXa, so it may exert cooperative effects on these regions. L2 is predicted to interact with the A2 domain. L3 is part of the basic heparin binding site of fXa [21] and is predicted to bind to the acidic portion of the a2-loop. L1 is composed of 14 residues, 5 of which are conserved, spanning from 124A to 135; L2 is 10 residues, of which 5 are conserved, spanning from 166 to 175; and L3 is 14 residues, of which 4 are conserved, spanning from 232 to 245 (Figure 1C). The effect of individual SP loop swaps on affinity for fV_BD_ was assessed by BLI, as above (Figure 3). All loop swaps improved affinity relative to wild-type fXa (Kd of 1375 nM), with L1 accounting for a 2.6-fold (Kd of 532 nM), L2 a 1.2-fold (Kd of 1134 nM), and L3 a 1.5-fold (Kd of 905 nM) decrease in Kd. Combining the three regions (L123) did not improve affinity beyond the effect of L1 on its own (Kd of 608 nM), and the L23 chimera was no better than either individual loop-swap (Kd of 1120 nM; Figure 3B). Importantly, combining the L123 chimera with the EGF2 swap yielded the ~15-fold increase in affinity seen for the whole SP domain swap (Kd of 608 to 43 nM; Figure 3C). However, when EGF2 was added to the L23 chimera, affinity only improved by about 3.7-fold (Kd of 1120 nM to 369 nM), suggesting cooperativity between L1 and EGF2. The L123/EGF2 chimera bound to fV_BD_ with a Kd of 43 nM, representing 88% of the binding energy gained from swapping the entire SP and EGF2 domains. This chimera is comprised of total of 45 mutations (M45) across three loops in the SP domain and the entire EGF2 domain (Figure 1C).

**Figure 3.**
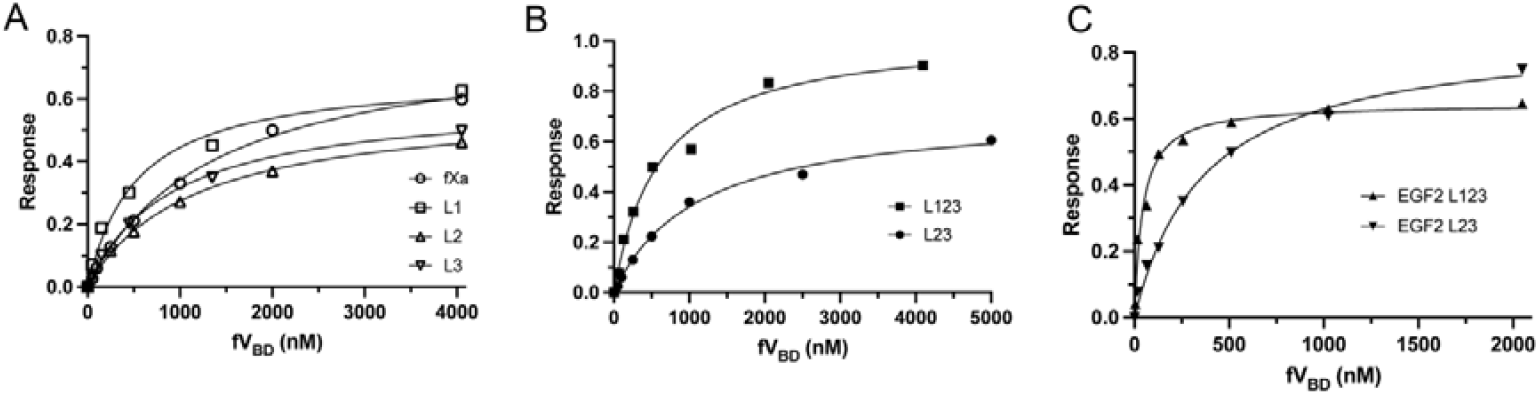
Binding of fXa loop-swap chimeras to fV_BD_ monitored by bio-layer interferometry (BLI). BLI response plotted against concentration of fV_BD_ is fit to a single site binding model to provide an estimate of the dissociation constant (Kd) for fXa and individual loop swap variants (L1, L2 and L3) as indicated (A), combined loop swap variants L123 and L23 (B), and L123 and L23 combined with the full EGF2 domain swap (C).

### M17 binds to fV_BD_ with high affinity

In order to reduce the number of mutations to just the key residues, we again examined the human prothrombinase model [13] to select only residues predicted to be making contact with fVa. We did not expect the model to be sufficiently accurate to predict actual side chain interactions, so this approach remained agnostic with respect to whether mutations were predicted to favor or disfavor binding. Seven of the original 21 mutations in the EGF2 domain were chosen based on this criterion, and in the SP domain, 4 of 9 mutations were selected from L1, 4 of 5 from L2, and 2 of 10 from L3 (Figure 1D). The resulting chimera had 17 mutations across the two domains and was denoted ‘M17’. Reverting to the wild-type sequence in L1 inadvertently introduced a consensus glycosylation sequence (NEV to NET; Figure 1C and 1D), but it was found to be unglycosylated in purified material (Supplemental Figure 2). As for the loop swap chimeras, we also made variants with subsets of mutations, including: M7 which only contains the 7 EGF2 mutations; and M13 which is missing the 4 L1 mutations. The affinities of M7, M13 and M17 for fV_BD_ were then assessed by BLI, as above. Once again, EGF2 was found to play a surprisingly large role in the interaction, with M7 binding with a Kd of 78 nM (Figure 4A). The addition of the 6 L2 and L3 mutations to M7 (M13) had a modest effect (Kd of 48 nM; Figure 4B). The M17 chimera was found to bind with high affinity, indistinguishable from the full domain-swap chimeras, with a Kd of 17 nM (Figure 4C). The EGF2 mutations therefore contribute 66% of the binding free energy difference, the L2 and L3 mutations an additional 9%, and the remaining 25% is contributed by the L1 mutations.

**Figure 4.**
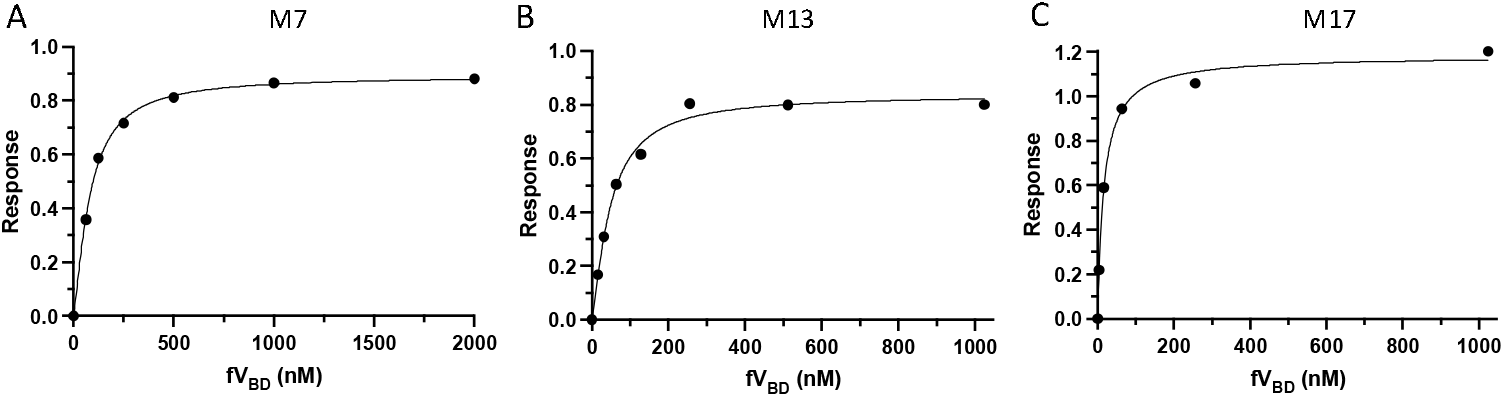
Binding of fXa variants to fV_BD_ monitored by bio-layer interferometry (BLI). BLI response plotted against concentration of fV_BD_ is fit to a single site binding model to provide an estimate of the dissociation constant (Kd) for fXa variants M7 (7 EGF2 mutations) (A), M13 (7 EGF2 and 6 SP mutations) (B) and M17 (C).

### Prothrombin processing by M17 fXa

We previously showed that HopD with fVa processes prothrombin exclusively down the meizothrombin pathway in the absence of PL membranes, similar to human prothrombinase with PL [12]. However, HopD prothrombinase lacks the processivity of human prothrombinase and gets stalled after cleavage of Arg320. In contrast, we found that prothrombin processing by M17 in the presence of fV_BD_ is rapid and complete (Figure 5A). The rates of prothrombin (II) consumption and the appearance of the heavy chain (H) are nearly identical (0.187±0.006 and 0.184±0.045 min^−1^, respectively), consistent with processing primarily down the meizothrombin route (Figure 5B). Interestingly, meizothrombin (F1.2-L) does not accumulate, with rates of its appearance and disappearance indistinguishable (both 0. 23±0.05 min^−1^, Figure 5C). This is similar to what we reported for human prothrombinase in the presence of PL [12] and is consistent with processive processing at Arg320 followed by Arg271. As with human prothrombinase, a small amount of prethrombin-2 (Pre-2) is evident, likely reflecting the activity of a fraction of uncomplexed fXa.

**Figure 5.**
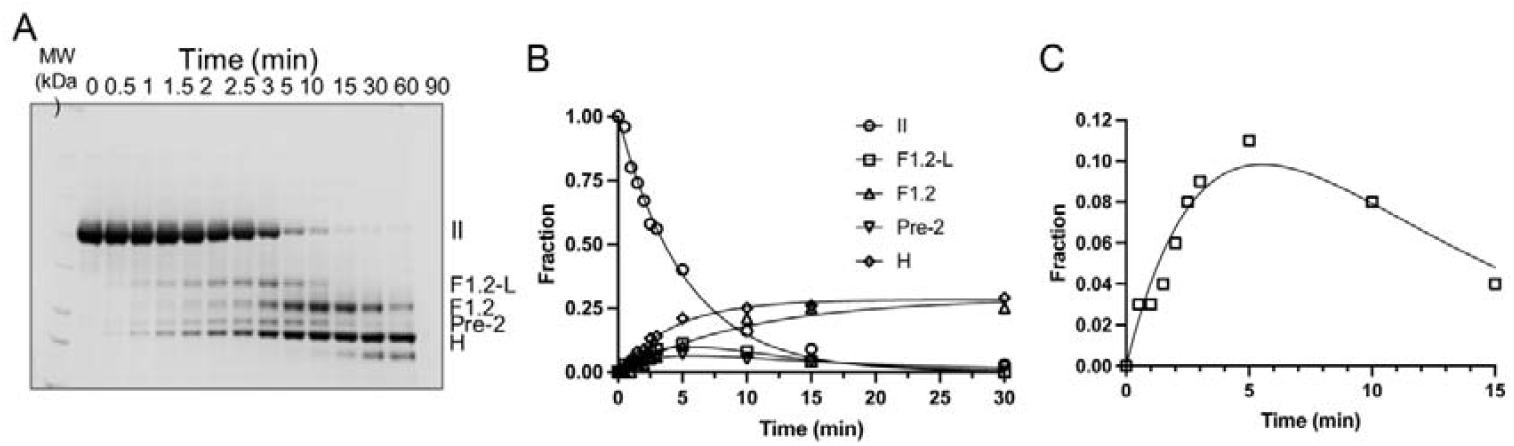
Prothrombin processing monitored by SDS-PAGE and analysis of cleavage products. (A) Cleavage of prothrombin (II) was monitored over a 90 min period by visualizing products on SDS-PAGE for M17 fXa with fV_BD_. Bands reflect cleavage events, with F1.2-L and H formed by cleavage at Arg320, and F1.2 and Pre-2 formed by cleavage at Arg271. The presence of F1.2-L indicates initial cleavage at Arg320, and the presence of Pre-2 indicates initial cleavage at Arg271. (B) Normalized densitometry of all bands plotted against time allows an evaluation of the kinetics of prothrombin processing. The solid lines reflect fits of II (circles) elimination and formation of the end products, H (diamond) and F1.2 (up triangle). Formation and elimination of intermediates Pre-2 (down triangle) and F1.2-L (square) are also fit. (C) A close up of the formation and elimination of F1.2-L is shown with fit yielding indistinguishable rates (a single representative experiment is shown).

In order to determine which regions were responsible for conferring processing down the meizothrombin pathway, we also assessed prothrombin processing with wild-type fXa and the L123 and EGF2-L123 chimeras in the presence of fV_BD_ without PL (Supplemental Figure 3). As expected, there was no evidence of meizothrombin formation for fXa, while increasing amounts of F1.2-L (the band reflecting initial cleavage at Arg320) were observed for L123 and EGF2-L123 (Supplemental Figure 4). The fraction of prothrombin processed via meizothrombin appears to be related to the affinity of fXa for fVa, and not to the region mutated in altering the affinity.

### Predicted effects of individual M17 mutations

To rationalize the effect of the M17 mutations on the improved binding to fVa, we conducted a close examination of each position and the nature of the substitution using our prothrombinase model and the recently published cryo-EM structure of prothrombinase (7TPP) [22], which was not available when this study was initiated. Superficially, our model and 7TPP look quite similar (Supplemental Figure 5A and B), however, they are substantially different, with an RMSD of 8.65 Å for 1,458 Cα atoms. When the fVa components are superimposed (Figure 6), it is clear that fXa is oriented differently in the two complexes, with fXa from 7TPP shifted by 8.8 Å and rotated by ~15° relative to its position in the model (using approximate center of mass of the heavy chain of fXa). The effect of the shift is to move fXa away from the body of fVa to accommodate the positioning of the a2-loop in 7TPP. In our model, the a2-loop protrudes straight out of the A2 domain to interact with the heparin binding site of fXa (Figure 6B), as observed in the crystal structure of pseutarin C [14]. In striking contrast, the a2-loop in 7TPP turns sharply across the A2 domain of fVa and mediates the bulk of the interactions with fXa [23]. The different position of fXa on fVa in 7TPP has the effect of placing most of the M17 mutations out of potential interaction distance with fVa.

**Figure 6.**
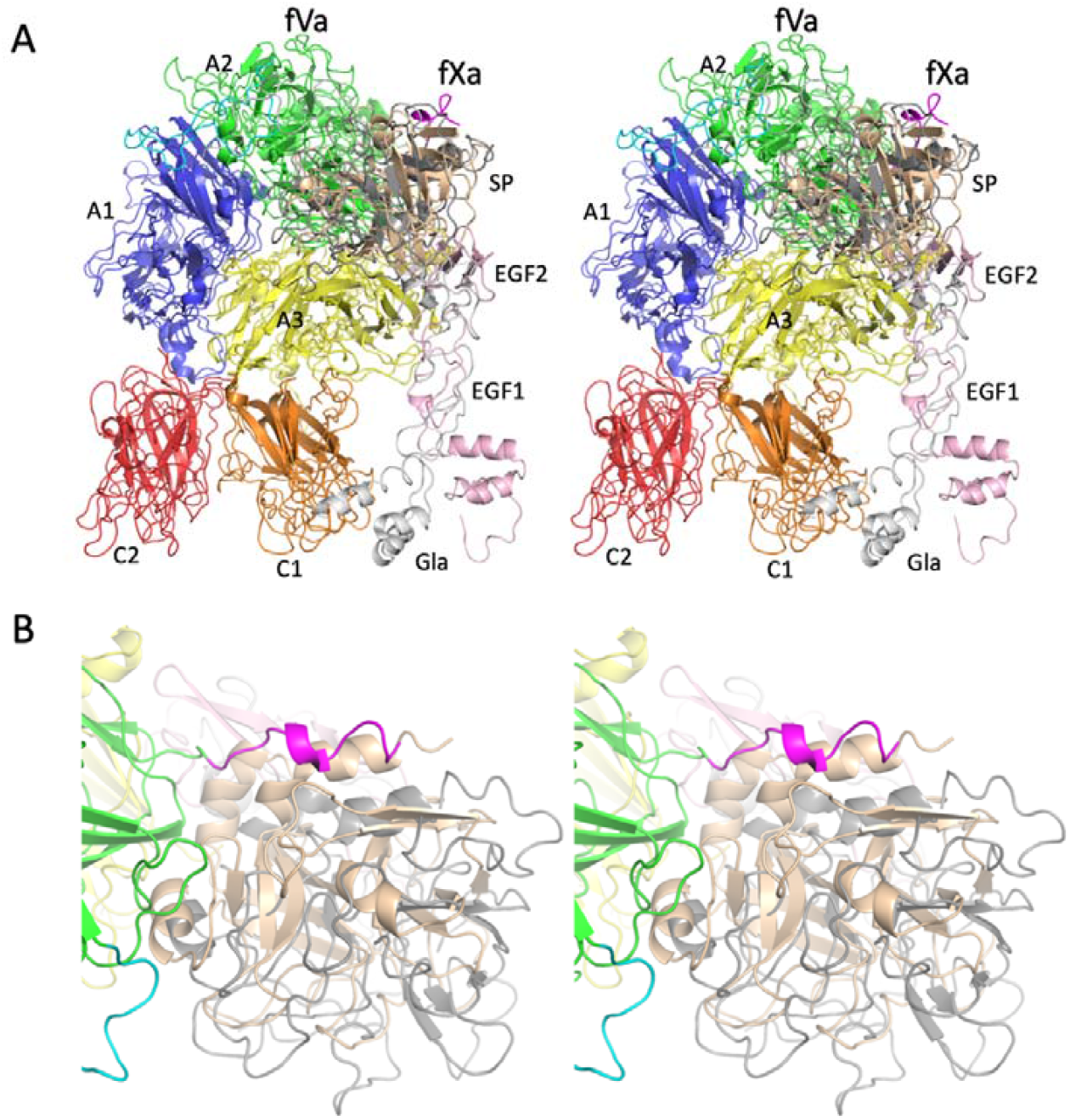
Stereo views of ribbon diagrams of a prothrombinase model and a cryo-EM structure (7TPP). (A) The model and cryo-EM structure are oriented identically, with fVa domains superimposed and colored as follows: A1 blue, A2 green, A3 yellow, C1 orange and C2 red, with the loop between the A1 and A2 domains cyan and the stretch C-terminal to the A2 domain in the model (the ‘a2-loop’) magenta. The SP domain of fXa is colored wheat for the model and dark gray for 7TPP, and the light chains are light pink (model) and light gray (7TPP). (B) A close-up of the SP domain is shown to illustrate the shift and rotation of the fXa component in 7TPP relative to the model (colored as in A). Figures were made using Pymol [32].

The predicted effect of each of the 17 mutations in M17 was assessed for our model and 7TPP by examining possible interactions (Supplemental Figures 6 and 7) and scoring the substitution as: ** when the substitution is clearly favored over the original; * when either the original or the substitution is predicted to contribute positively to binding; ^^ when the mutated residue is unable to make any contact with fVa (at too great a distance); and, ^ where the original residue is likely to be better than the substitution. This scoring system was then quantified by assigning values of ** = +2, * = +1, ^^ = −1 and ^ = −1, and the results of this analysis are given in Supplemental Tables 1 and 2. Bearing in mind that the 17 mutations improve affinity of fXa for fVa by ~50-fold, it is reasonable to expect that most of the substitutions should be at the interface and making favorable contacts. In our model, 7 of the mutations were ranked as better than the original (**) and 4 as making good contacts as either original or mutation (*). In 4 positions the original residues are predicted to be better than the substitution (^), and in 2 the mutated residues are too distant from fVa to make any contact (^^). The total score is +12. Interestingly, +6 is contributed by mutations in the EGF2 domain, +4 by L1, and only +1 by each L2 and L3, similar to the contribution of each of these regions to improved binding affinity. In contrast, analysis of the predicted effect of the M17 mutations on the structure 7TPP yields a score of −10, with only 1 substitution predicted to be better than the original, 2 as either might work, 5 as being better in the original, and 9 as not predicted to make any contact. If we score residues outside contact range as a 0 instead of −1, the model receives a score of +18 and 7TPP is still negative at −1. Although the model is unlikely to be correct in detail, it is clear that 7TPP is inconsistent with the M17 mutations having the observed effect of improved affinity for fVa.

## DISCUSSION

The membrane dependence of prothrombinase assembly and function ensures that thrombin generation is limited to sites of vascular damage where phosphatidylserine (PS)-rich PL surfaces are expressed. The importance of PL membranes in the regulation of prothrombinase is illustrated by the prothrombinase orthologues found in the venom of snakes of the Elapidae family in Eastern Australia [11]. The fXa orthologues from the venom of group D snakes, such as HopD, are lethal when injected into rodents causing disseminated intravascular coagulation, organ failure and hemorrhage [24,25]. The toxicity of HopD is conferred by its ability to bind to fVa with high affinity in the absence of PL membranes [12]. We set out to create a similar membrane independent human fXa by determining the contributions of individual domains to binding affinity, and then narrowing the scope of the substitutions to regions predicted to be at the interface with fVa.

With just 17 mutations across the EGF2 and SP domains of fXa we were able to recapitulate the high affinity of HopD for fVa, and importantly, to switch the processing of prothrombin so that it proceeds via the meizothrombin pathway in the absence of PL membranes. Interestingly, unlike HopD which stalls after initial cleavage of Arg320, cleavage of meizothrombin at Arg271 by M17 in the presence of fVa is rapid and kinetically indistinguishable from meizothrombin formation (initial cleavage at Arg320), the hallmark of processivity. The processivity observed for human prothrombinase has been well documented and is best explained by the ‘ratcheting’ mechanism [26], where cleavage of Arg320 promotes a conformational rearrangement of the SP domain and its interface with the second kringle domain to reposition meizothrombin on fVa [27], thereby presenting Arg271 to the active site of fXa without requiring meizothrombin to dissociate. Although dissociation of the intermediate meizothrombin can and evidently does occur in vitro, since some intermediate can be observed by SDS-PAGE, the inability to distinguish between the rates of Arg320 and Arg271 cleavage suggests that dissociation between cleavage events is rare or that re-association is fast [28].

The accumulation of meizothrombin using HopD-prothrombinase suggests that dissociation of meizothrombin is faster than the presentation of Arg271 to the active site of HopD. Meizothrombin would then need to compete with prothrombin for re-association to HopD-prothrombinase for the second cleavage event to occur-thus breaking the ratchetting mechanism. Since the fVa component is the same whether HopD or M17 is bound, it is logical to conclude that the fXa component is the reason for the differences in prothrombin processing. Testing this hypothesis is reasonably straightforward either using a chimeric approach to swap loops from HopD into fXa that are likely to be mediating contact with prothrombin.

Although the affinity of M17 for fVa is similar to that of HopD for fVa, M17-prothrombinase appears to use both pathways, with evidence of the Pre-2 intermediate, whereas HopD-prothrombinase processes prothrombin exclusively through the meizothrombin intermediate. The presence of two insertion loops on the SP domain of HopD relative to fXa may be partially responsible for this difference. In addition, HopD contains a Glu at position 192, in place of Gln found in all other fXa orthologues, which reduces its ability to cleave prothrombin when not in complex with fVa [12]. There is therefore no leaky processing via Arg271 by uncomplexed HopD as there is for fXa/M17, which can be misinterpreted as an alternate processing route for assembled prothrombinase. If the insertion loops or other sequence differences at the prothrombin binding interface of HopD enforce initial cleavage at Arg320, then it is unlikely that improving the affinity of human fXa for fVa in the absence of PL would be sufficient to confer processing down the meizothrombin pathway. If, on the other hand, the observed processing via Pre-2 for human fXa under any condition simply reflects the fraction of uncomplexed fXa, then improving its affinity for fVa will reduce flux down the Pre-2 pathway in favor of the meizothrombin pathway.

Although the results presented here do not rule out the possibility of alternate processing by assembled prothrombinase, they strongly support the model where processing of prothrombin via Pre-2 only occurs when fXa is not in complex with fVa. Transient intermediates, especially ones that are processivity processed like meizothrombin, are hard to detect. However, we were able to observe a band corresponding to initial cleavage at Arg320 (F1.2-L) in the absence of membranes after improving the affinity of wild-type fXa by only 2.3-fold (Kd from 1375 nM to 608 nM; Supplemental Figure 3). The fraction processed via the meizothrombin route increased with each improvement in affinity and the fraction of Pre-2 decreased. However, processing of prothrombin by our best variant, M17, which has a Kd for fVa similar to that of HopD, still results in a band corresponding to the intermediate Pre-2 by SDS-PAGE. On the surface this appears to argue against the ‘leaky’ processing hypothesis. However, M17 binds with a Kd of ~20 nM, so under the conditions used to monitor prothrombin processing (5 nM M17 and 50 nM fV_BD_) only about 70% of the M17 is complexed with 30% free to make Pre-2. And since processing of prothrombin (5 μM initial concentration) is likely to be much faster than processing of the Pre-2 product by either free or complexed M17, Pre-2 will accumulate. Experiments of this nature are therefore skewed towards observing Pre-2 as an intermediate, whilst the processivity of the meizothrombin intermediate biases against its accumulation.

It is clear that the PL surface improves the affinity of fXa for fVa, and that assembly is required for efficient processing of prothrombin. However, it is believed that prothrombinase assembly on its own is insufficient to confer processing via meizothrombin, and that the interaction between the substrate prothrombin and the PL surface is required to enforce the order of cleavage. Two reports describe the processing of prothrombin lacking the post-translational modification of gamma carboxylation [9,10], which is therefore unable to bind to Ca^2+^ ions and interact with PS-containing membranes with high affinity. Both studies, one using bovine and the other human proteins, found that processing occurred preferentially down the Pre-2 pathway, with no evidence on SDS-PAGE of the meizothrombin intermediate. The authors of the more recent report concluded that membrane tethering is essential in ‘dictating the presentation of the substrate to prothrombinase’ [9]. Our results appear to contradict these findings by suggesting that the membrane is irrelevant in the presentation of Arg320 for initial cleavage, provided prothrombinase is assembled. Our findings are consistent with earlier work on bovine prothrombinase that inferred from kinetics that processing occurs substantially via meizothrombin when fVa is saturating in the absence of PL [29].

This apparent discrepancy may be due to the use of un-gamma carboxylated prothrombin and 25% PS vesicles. Without gamma carboxylation, the Gla domain will be unable to bind Ca^2+^ and will be in a non-native, artifactual conformation, and with no divalent cations to counterbalance the high negative charge. Comparison of the AlphaFold [30] model of human des-Gla F1 with the crystal structure of gamma carboxylated bovine F1 [31] bound to Ca^2+^ (Supplemental Figure 8) reveals major differences in the electrostatic surface presented to the PL membrane. This ‘des-Gla’ prothrombin might be repulsed from the highly negatively charged surface of the PS-rich vesicles, inhibiting the K2-SP domains from interacting with fVa in the normal fashion. This explanation is supported by work on a des-Gla form of prothrombin generated by chymotrypsin treatment which showed no difference in rate of thrombin generation when PL was added to a fVa titration [4]. This experiment measured only thrombin activity, so effectively monitored cleavage at Arg320. Since PL made no difference to the maximal rate, it is unlikely that PL is involved in presentation of Arg320 to the active site of fXa.

It is interesting to note that the improved affinity of M17 for fVa appears to be inconsistent with the recently published cryo-EM structure of prothrombinase (7TPP) [22]. This is likely due to the poor quality and low resolution of the map used to build the model [23]. A low resolution cryo-EM map provides little more than an envelope into which proteins and domains of known structure can be placed. The degree of accuracy of the placement depends on the information content of the map. The map for 7TPP is of exceedingly low resolution (likely 8-10 Å, instead of the claimed 4.1 Å) and its information content is further limited by an extreme orientation bias of the particles used to calculate the map. The inconsistency of the M17 mutations with 7TPP supports further efforts to obtain a detailed structure of the prothrombinase complex, perhaps with the aid of a high-affinity variant of fXa such as M17.

## Authorship Contributions

JAH and FIU conceived of and designed the study. FIU conducted the experiments. JAH and FIU interpreted the data, and JAH wrote the manuscript.

