## Supplemental Information for "Engineering a membrane-independent human prothrombinase through parsimonious mutation of factor Xa"

**Supplemental Table 1.** M17 substitutions scored as described in the main text for the model of prothrombinase based on the crystal structure of pseutarin C, with comments.

| | substitution | score | comments on contacts with fVa |
|---|---|---|---|
| EGF2 | S90R | ** | near D628 and E1650 |
| | L91A | * | in hydrophobic patch next to V1627 and V1681 |
| | D92F | ^ | D is in contact with N-term of light chain N1547; no reason to expect an F would be better |
| | H101K | ^^ | H is not in contact with anything, K will have no effect |
| | E102R | ** | in a hydrophobic area near F1666 and E1650; R could pick up hydrophobic and electrostatic interactions |
| | E103V | ** | in hydrophobic area near P1663, W1665 and F1666; V definitely better |
| | N105S | * | hydrophobic area near W1665 and Y1678; S might fit better |
| L1 | E129N | ** | near D577, so N relieves electrostatic repulsion |
| | S130E | ** | in hydrophobic site near F576 and V630, but very close to K655 |
| | T132K | * | near D1625 but also near H1683; unclear |
| | K134D | ^ | looks ideal as a K; makes electrostatic contact with D1625 and E1686; predicted negative effect of mutation to D |
| L2 | N166H | * | hydrophobic region near A511, I604 and T624; no reason to expect H would be better |
| | K169M | ^ | looks optimal for a K; near D513, D577 and D578; not hydrophobic at all; predict K is much better |
| | S173D | ** | right next to R317, so D would be much better; S is making no side-chain contact |
| | I175R | ^ | basic and hydrophobic region, near L503, R505, R510; change to R predicted to have negative effect |
| L3 | A233R | ** | near D578, D659 and D660; R would also pick up hydrophobic interactions with I657 and P658 |
| | D239K | ^^ | no predicted interactions; on wrong side of C-terminal helix to contact the a2-loop |

**Supplemental Table 2.** M17 substitutions scored as described in the main text for the cryo-EM structure of prothrombinase from 7TPP, with comments.

| | substitution | score | comments on contacts with fVa |
|---|---|---|---|
| EGF2 | S90R | ** | far from fVa, but might pick up int with E1650 |
| | L91A | * | in contact with H1683; A will make no difference; no favorable contacts for either |
| | D92F | ^ | right up against D1625; mutation to F would not be favored; not a hydrophobic area |
| | H101K | ^^ | too far from fVa to make any contact |
| | E102R | ^^ | too far from fVa to make any contact |
| | E103V | ^^ | too far from fVa to make any contact |
| | N105S | ^^ | too far from fVa to make any contact |
| L1 | E129N | ^^ | too far from fVa to make any contact |
| | S130E | ^^ | too far from fVa to make any contact |
| | T132K | ^^ | too far from fVa to make any contact |
| | K134D | ^^ | too far from fVa to make any contact |
| L2 | N166H | ^ | hydrophobic region with T579, I604, T626; making contact with T579; H would not be favorable |
| | K169M | ^ | not making any electrostatic contacts, but could do with D577; a hydrophilic area; no advantage of an M |
| | S173D | ^ | mutation would be very bad; right up against E691, and making close contact with E401 of fXa (E217 in chymotrypsin numbering) |
| | I175R | * | R could pick up interactions with D683, E690; but is right next to R510 |
| L3 | A233R | ^ | R would not fit; A is buried and tightly packed; very buried |
| | D239K | ^^ | too far from fVa to make any contact |

Key: ** if substitution is better than the original; * if both original and substitution are favorable; ^^ if residue makes no contact with fVa; ^ if original is better than the substitution.

**Supplemental Figure 1. Assessment of chromogenic activity of fXa chimeras (A and B) and variants (C) against the substrate S-2222.** To assess the effect of domain swaps and mutations on the activity of fXa variants towards a chromogenic substrate, proteins were diluted in TBS pH 7.4 buffer containing 5 mM $CaCl_2$ and added to 96-well plates (final concentration of 10 µg/ml in 50 µl). Reactions were initiated by adding 50µl of 0.4 mM S-2222 to each well. Absorbance values were monitored at 405 nm for 10 minutes using Versamax plate reader (Molecular Devices).

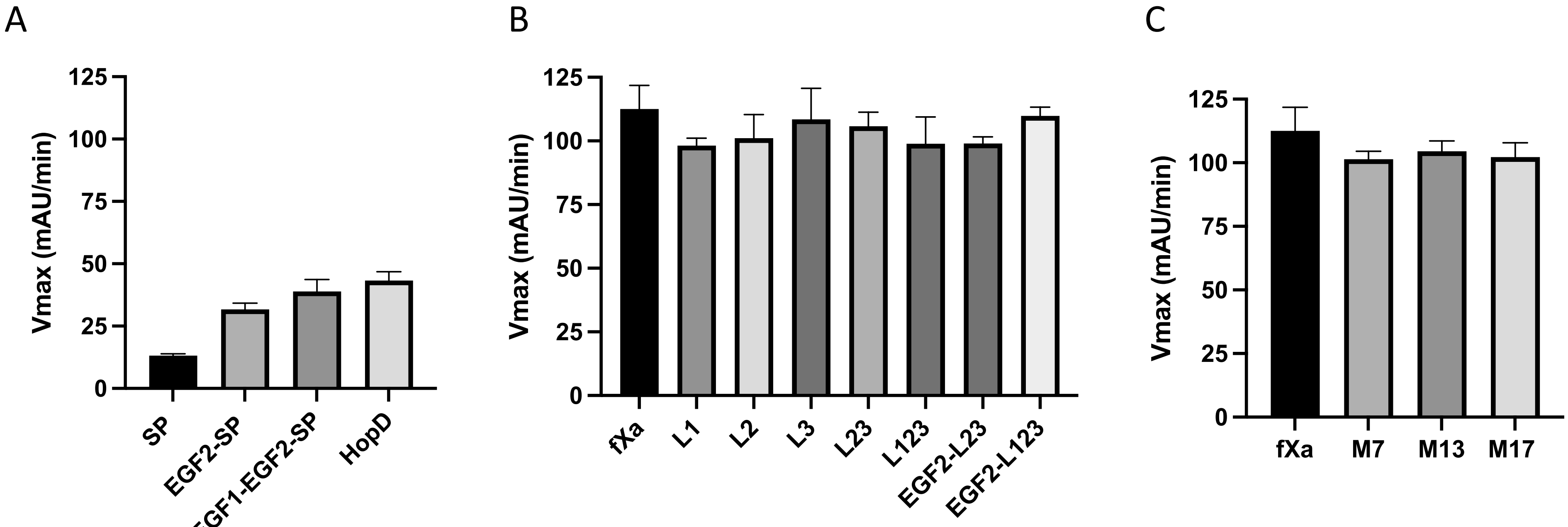

**Supplemental Figure 2. Assessment of M17 glycosylation.** To directly assess the fraction of glycosylated M17 we utilized a concanavalin A agarose resin kit. Wild-type fXa does not contain any glycosylation sites, so M17 should only bind to the column if a glycan is introduced. As a control, we tested E. coli and mammalian cell-derived thrombin (single glycosylation site at Asn60G in chymotrypsin numbering) and found that only the mammalian-derived thrombin bound to the resin and eluted upon treatment with elution buffer, while E. coli-derived material came off during the loading and wash steps (A). M17 eluted during the loading and wash steps, with no fXa evident on the gel in the lane corresponding to the elution fraction (B), nor was any S-2222 cleavage activity detected in the elution fraction (data not shown). Although a glycosylation consensus sequence was introduced in M17, no N-linked glycosylation is present in the material purified from our HEK-EBNA expression system.

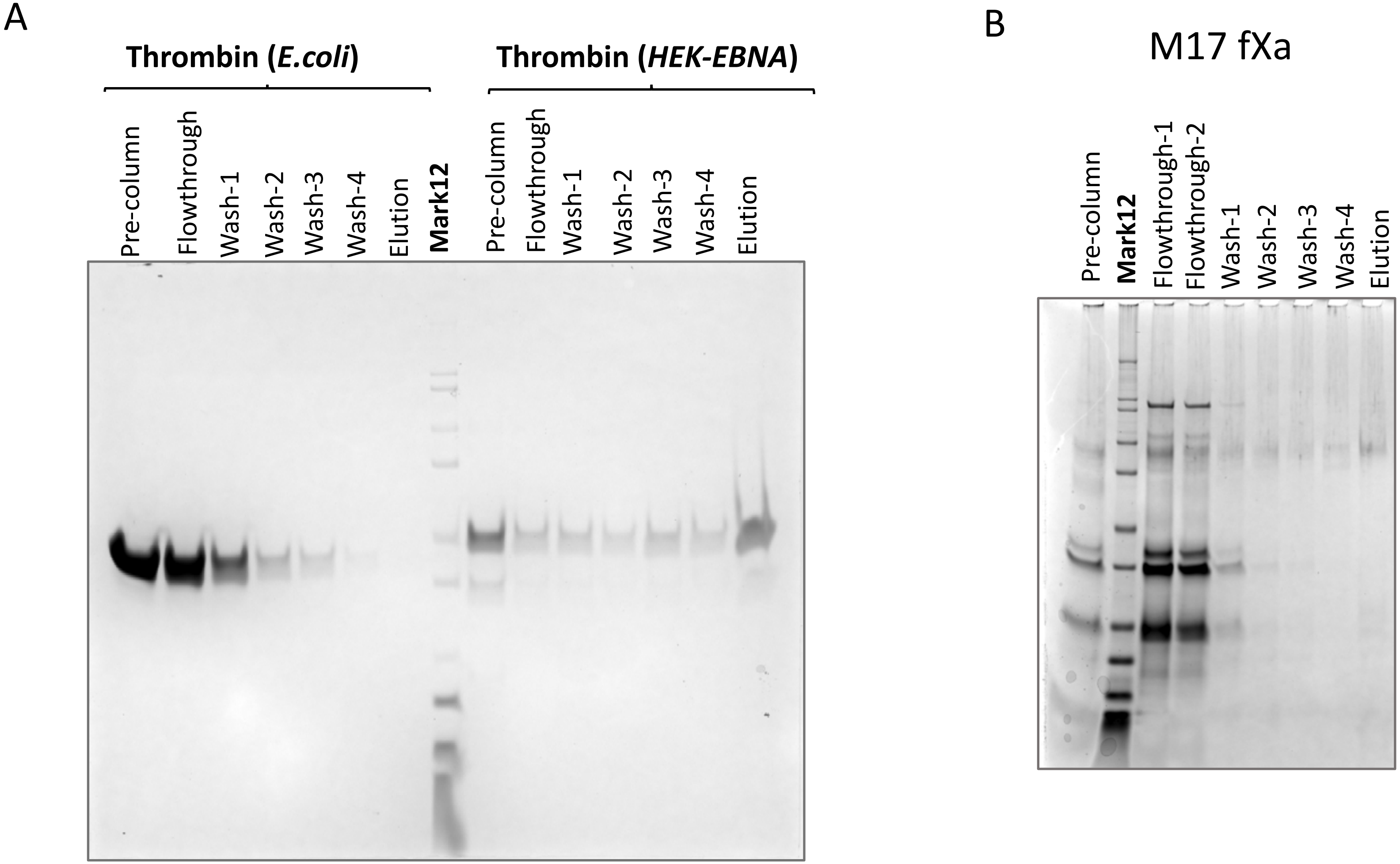

**Supplemental Figure 3. Prothrombin processing monitored by SDS-PAGE and analysis of cleavage products.** Cleavage of prothrombin (II) was monitored over a 90 min period by visualizing products on SDS-PAGE for wild-type (WT) fXa (A), L123 (B) and EGF2/L123 (C) chimeras with $fV_{BD}$ in the absence of PL membranes. Bands reflect cleavage events, with F1.2-L and H formed by cleavage at Arg320, and F1.2 and Pre-2 formed by cleavage at Arg271. The presence of F1.2-L indicates initial cleavage at Arg320, and the presence of Pre-2 indicates initial cleavage at Arg271. Plots of normalized densitometry of all bands plotted against time is shown in the lower panel. The solid lines represent fits of II (circles) elimination and formation of the end products, H (diamond) and F1.2 (up triangle). Formation and elimination of intermediates Pre-2 (down triangle) and F1.2-L (square) are also fit.

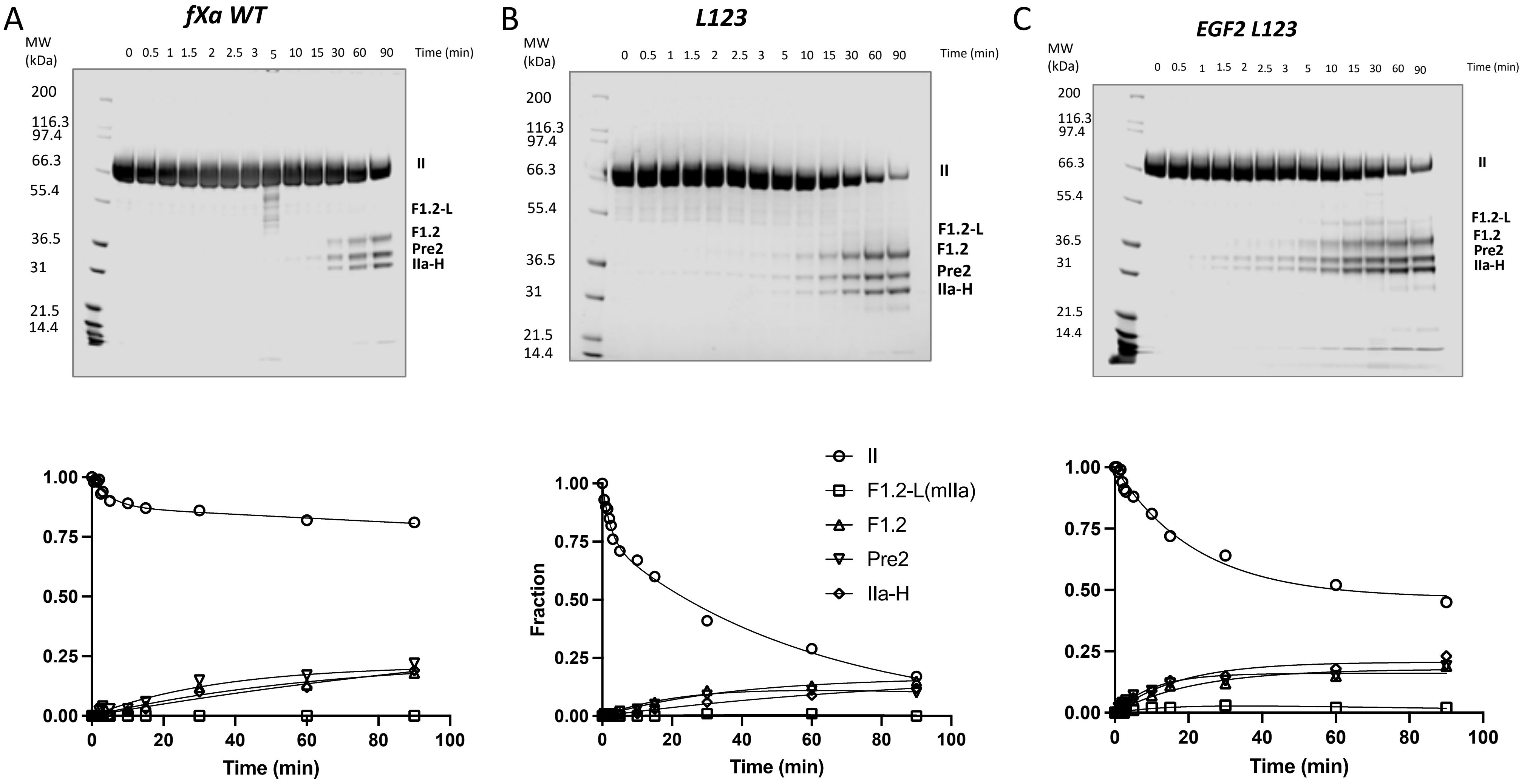

**Supplemental Figure 4. Appearance and disappearance of band corresponding to F1.2-L reflects processing via meizothrombin intermediate.** No detectable band for F1.2-L is observed for wild-type fXa (crosses), with very little evident for the L123 chimera (plus sign). The increase in affinity afforded by the addition of the EGF2 swap to the L123 chimera (circles) results in more processing through initial cleavage at Arg320 via the meizothrombin intermediate. As for HopD and M17, rates of formation and elimination are indistinguishable.

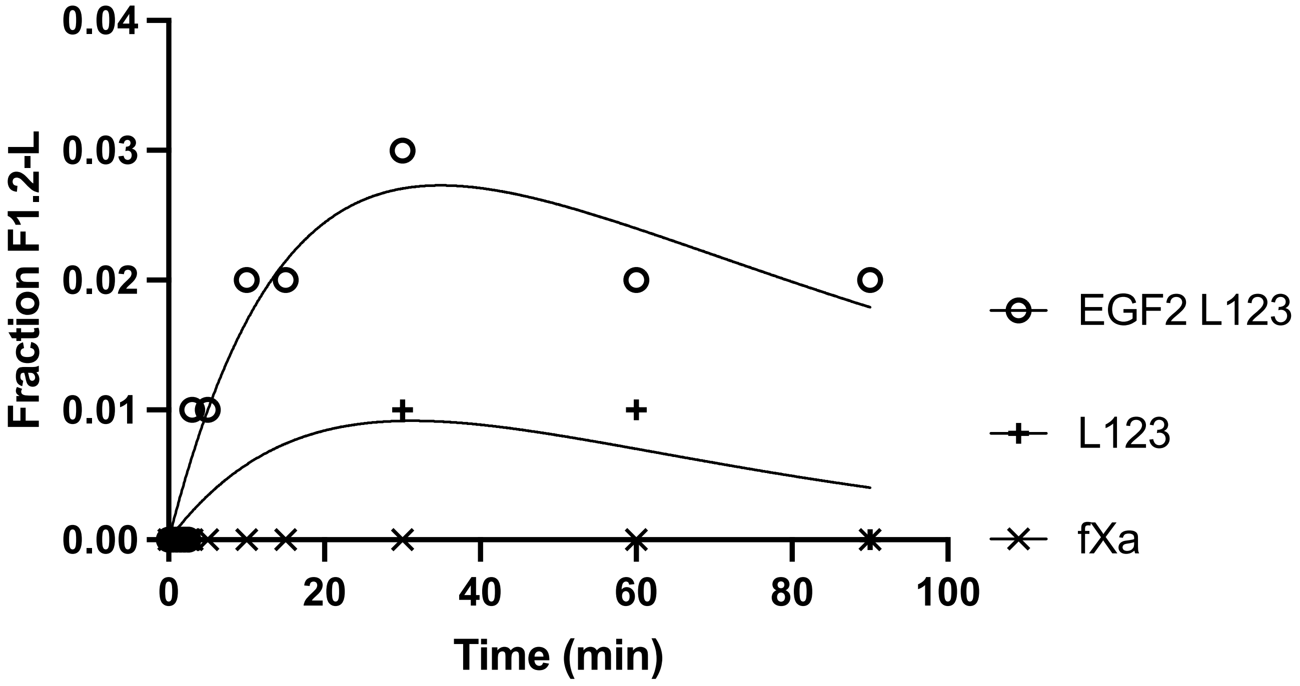

**Supplemental Figure 5. Stereo views of ribbon diagrams of a prothrombinase model (A) and a cryo-EM structure (B; 7TPP) illustrate a superficial similarity.** Both panels are oriented identically with fVa domains colored as follows: A1 blue, A2 green, A3 yellow, C1 orange and C2 red, with the loop between the A1 and A2 domains cyan and the a2-loop magenta (for model only). The fXa heavy chain is colored wheat for the model and dark gray for 7TPP, and the light chains are light pink (model) and light gray (7TPP).

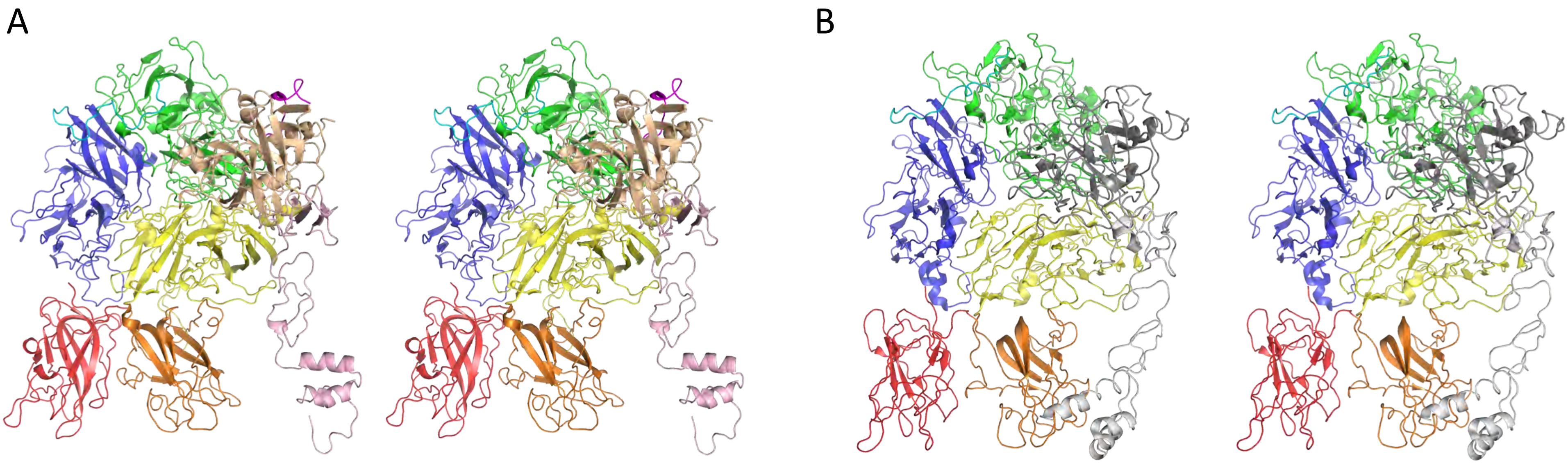

**Supplemental Figure 6. M17 residues and the prothrombinase model.** The M17 side chains are depicted as sticks, with all other sidechains in wire representation. The domain coloring is the same as in Figure 5 in the main manuscript and Supplemental Figure 4.

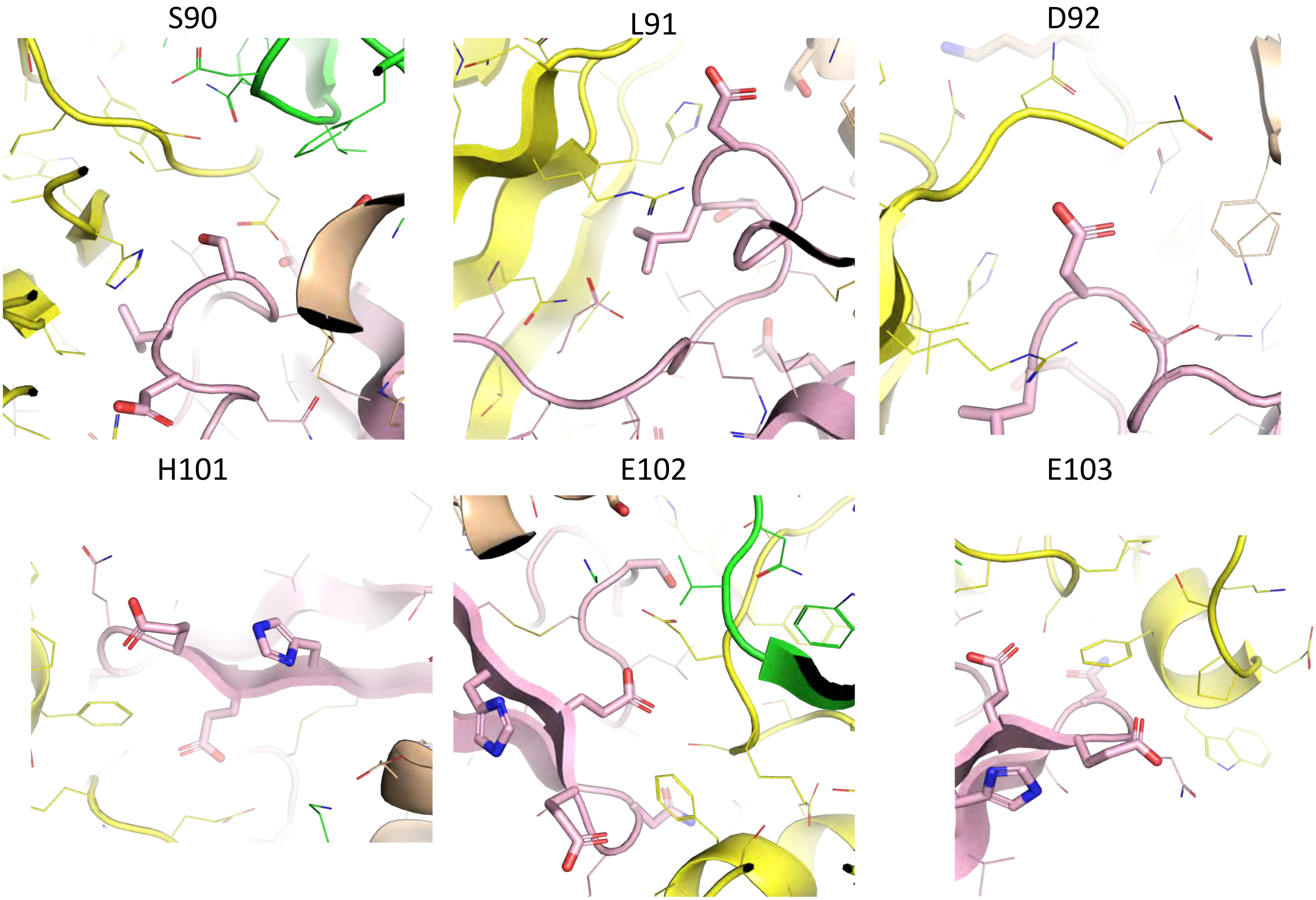

**Supplemental Figure 6, continued.**

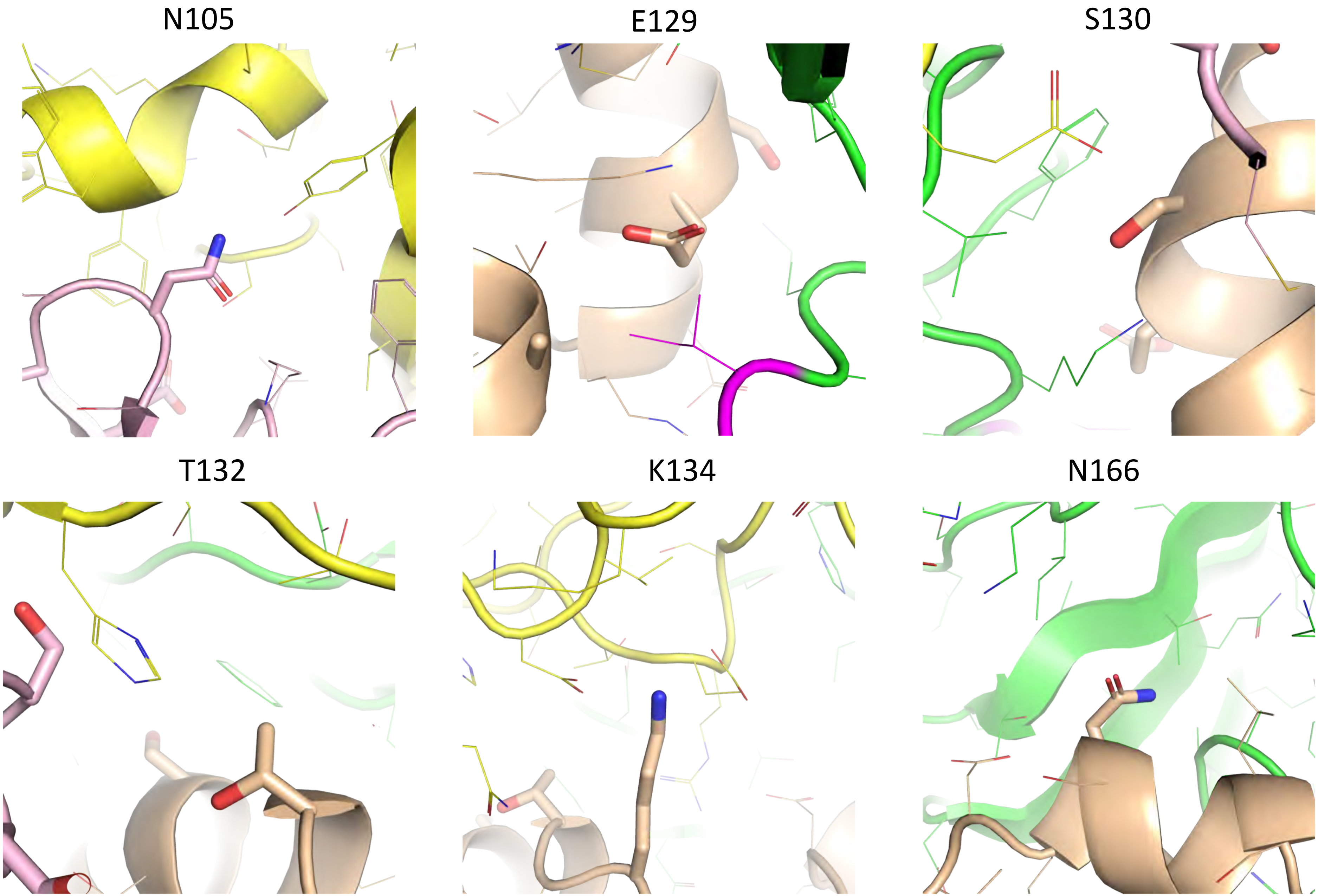

**Supplemental Figure 6, continued.**

K169

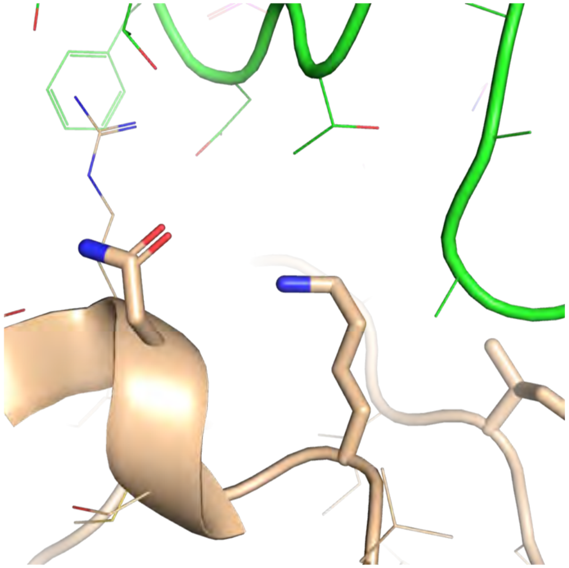

S173

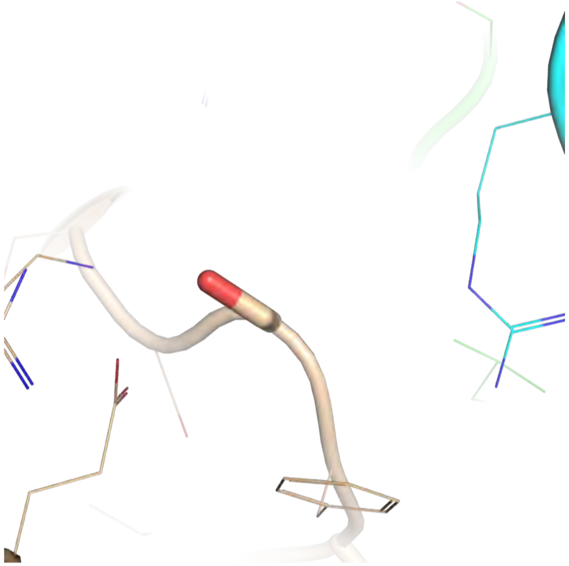

I175

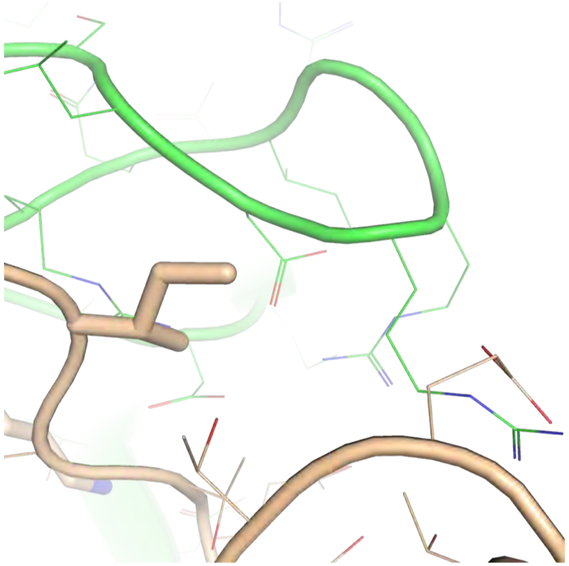

A233

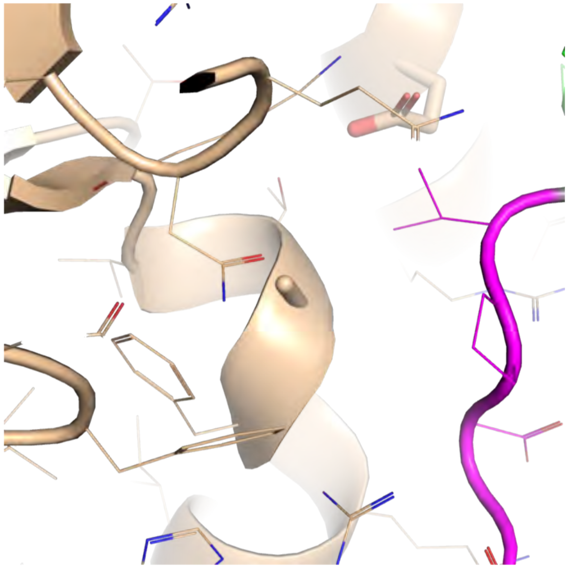

D239

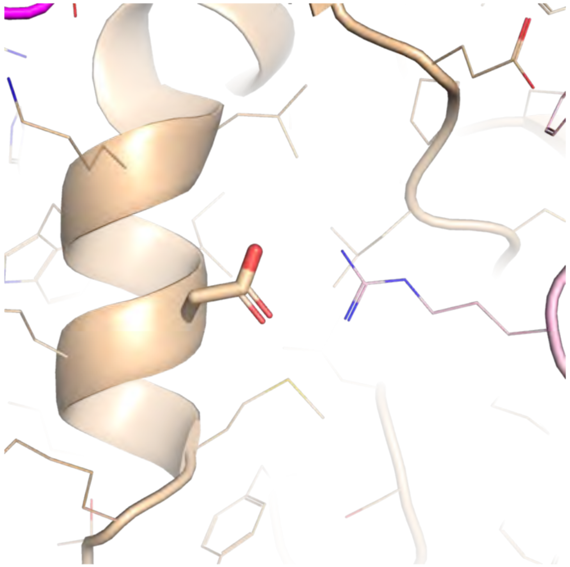

**Supplemental Figure 7. The M17 residues and prothrombinase from 7TPP.** The M17 side chains are depicted as sticks, with all other sidechains in wire representation. The domain coloring is the same as in Figure 5 in the main manuscript and Supplemental Figure 4.

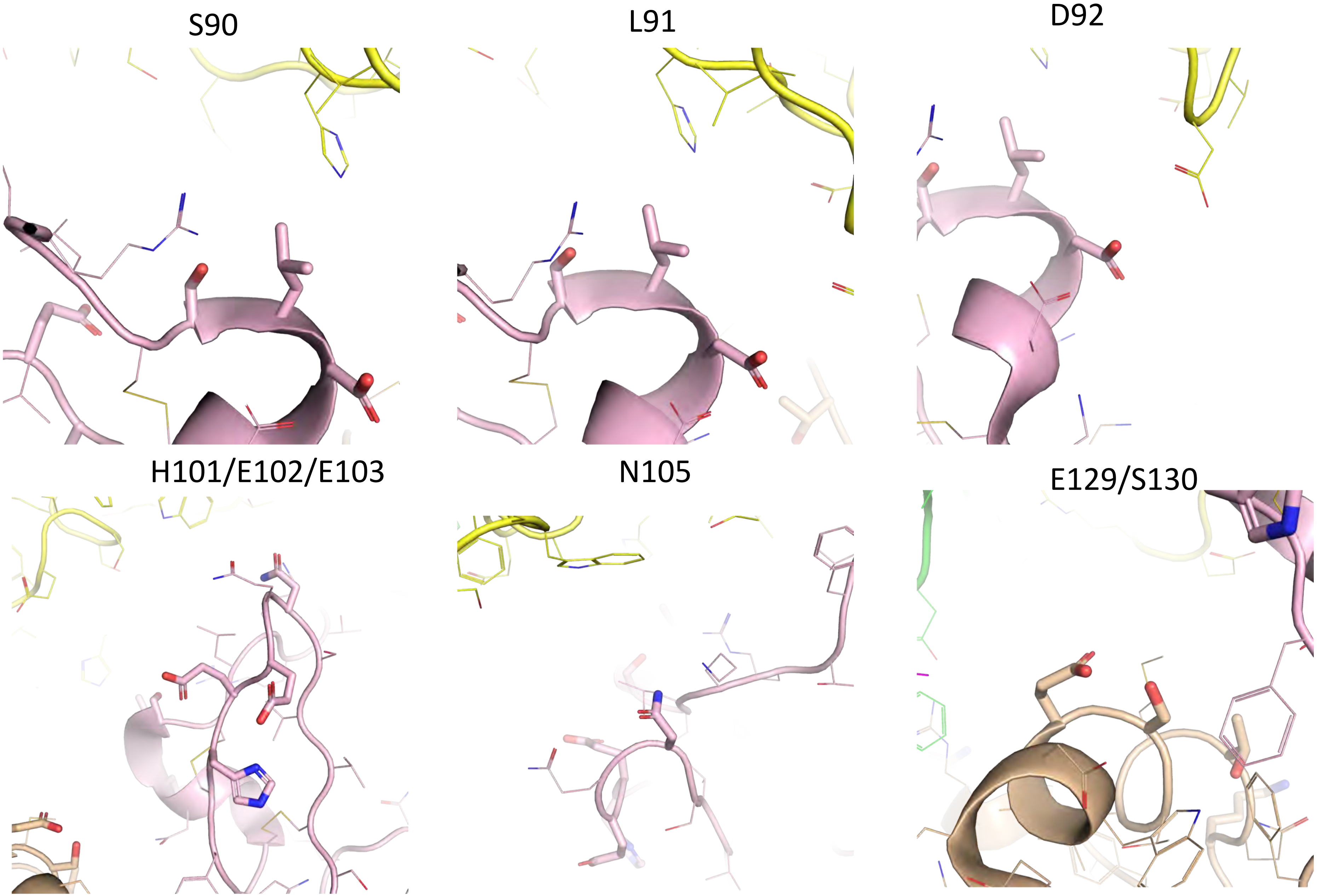

**Supplemental Figure 7, continued.**

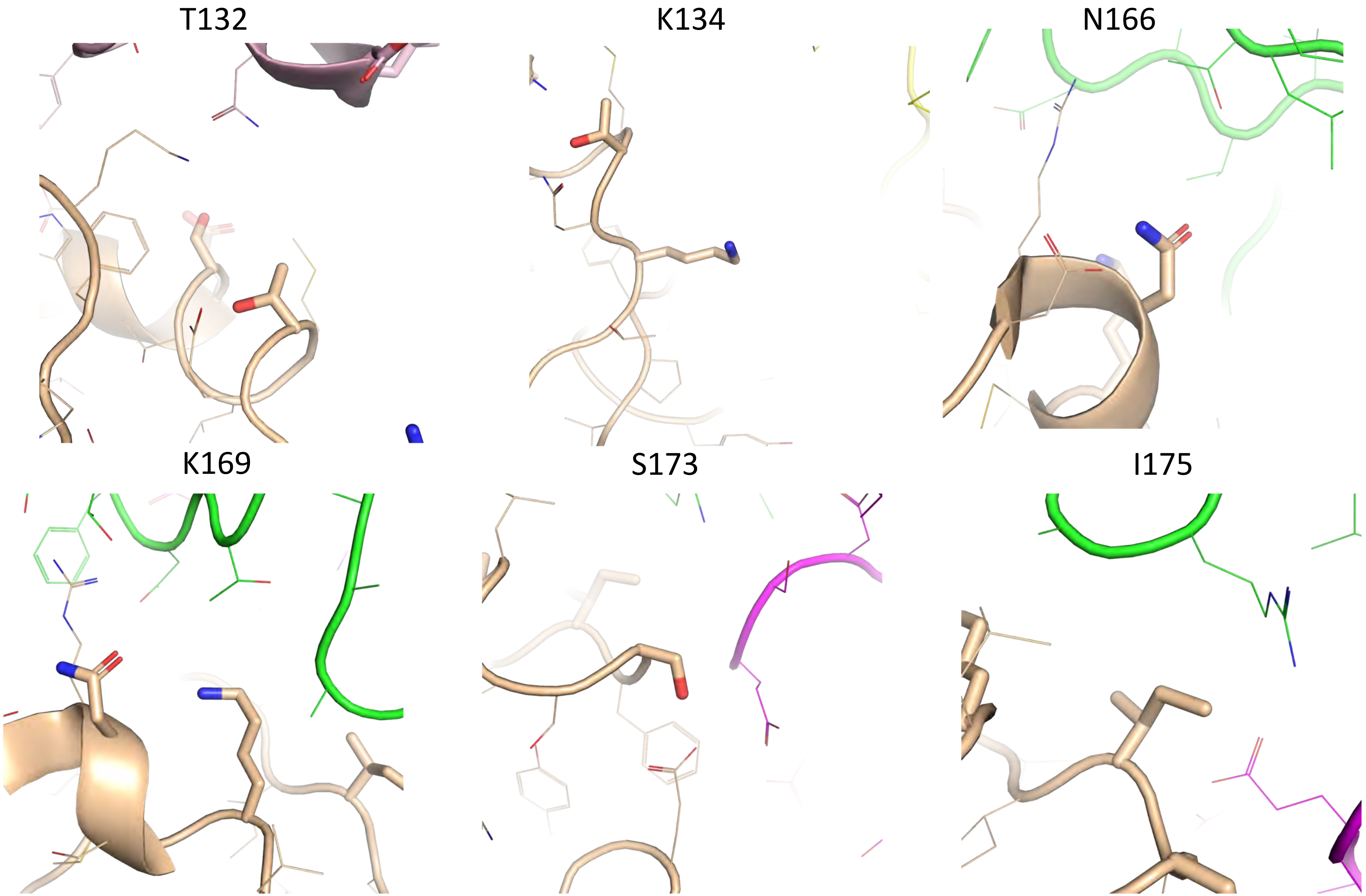

**Supplemental Figure 7, continued.**

A233

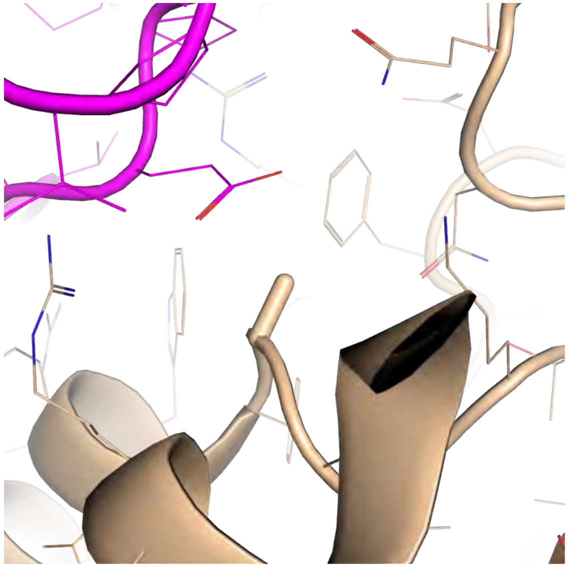

D239

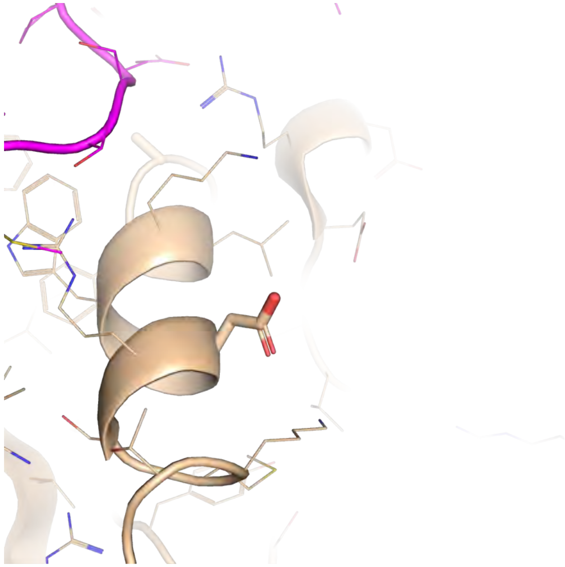

**Supplemental Figure 8. Stereo views of the F1 region of prothrombin (Gla-K1 domains) from the structure of bovine F1 (2PF2, cyan) with $Ca^{2+}$ (green balls) (A) and the AlphaFold predicted structure of human F1 without gamma carboxylation (B).** Cartoon representation is shown in the upper panel and electrostatic surface representation in the lower panel. View is of the bottom of the Gla domain, the region that should interact with the negatively charged phospholipid membrane.

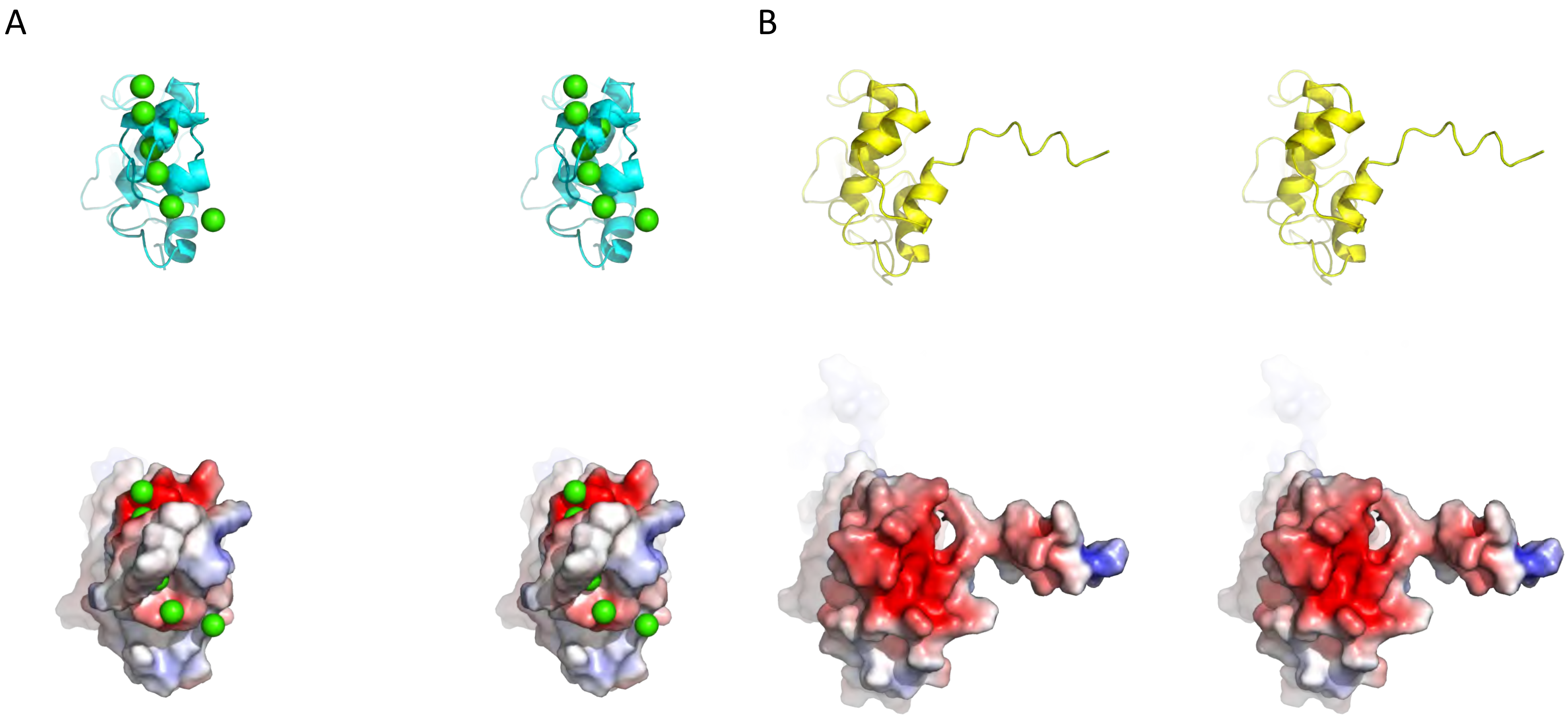
